# A Mechanism Based Pharmacokinetic/Pharmacodynamic Analysis of Polymyxin B-Based Combination Therapy Against Carbapenem-Resistant *Klebsiella pneumoniae* Isolates with Diverse Phenotypic and Genotypic Resistance Mechanisms

**DOI:** 10.1101/2025.05.06.652358

**Authors:** Ramya Mahadevan, Estefany Garcia, Rajnikant Sharma, Hongqiang Qiu, Ahmed Elsheikh, Robert Parambi, Cely Saad Abboud, Fernando Pasteran, Maria Soledad Ramirez, Keith S. Kaye, Robert A. Bonomo, Gauri G. Rao

**Affiliations:** Titus Family Department of Clinical Practice, USC Alfred E. Mann School of Pharmacy and Pharmaceutical Sciences, University of Southern California, Los Angeles, California, USA; Division of Pharmaceutics and Experimental Therapeutics, Eshelman School of Pharmacy, University of North Carolina at Chapel Hill, Chapel Hill, NC, USA; Department of Pharmacy, Fujian Medical University Union Hospital, Fuzhou, 350001, People’s Republic of China; College of Pharmacy, Fujian Medical University, Fuzhou, 350004, People’s Republic of China; Instituto Dante Pazzanese de Cardiologia, São Paulo, Brazil; Division of Allergy, Immunology and Infectious Diseases, Rutgers Robert Wood Johnson Medical School, New Brunswick, New Jersey, USA; Antimicrobianos, Instituto Nacional de Enfermedades Infecciosas, Antimicrobial Service of the National Institute of Infectious Diseases (ANLIS Dr. Carlos G. Malbrán), Buenos Aires, Argentina; Center for Applied Biotechnology Studies, Department of Biological Science, College of Natural Sciences and Mathematics, California State University Fullerton, Fullerton, CA, USA; Department of Molecular Biology and Microbiology, Case Western Reserve University School of Medicine, Cleveland, Ohio, USA; CWRU-Cleveland VAMC Center for Antimicrobial Resistance and Epidemiology (Case VA CARES), Cleveland, Ohio, USA; Geriatric Research, Education and Clinical Center, Louis Stokes Cleveland Department of Veterans Affairs Medical Center, Cleveland, Ohio, USA; Department of Medicine, Case Western Reserve University School of Medicine, Cleveland, Ohio, USA; Departments of Pharmacology, Biochemistry, and Proteomics and Bioinformatics, Case Western Reserve University School of Medicine, Cleveland, Ohio, USA

## Abstract

Increased resistance to β-lactams/β-lactamase inhibitor by mutations in β-lactamases gene, porin mutations and efflux pumps complicates the management of carbapenem-resistant *Klebsiella pneumoniae* (CRKP). Polymyxin B (PMB) based combination therapy is considered as a best alternative treatment for those middle and low-income countries that cannot access to the latest medicines. It’s crucial to know both phenotypic and genotypic characteristics of a pathogen to understand the killing effect of each drug and its combinations. Hence, our objective of this study was to incorporate mechanistic insights gained from resistance mechanisms to develop a mechanism based pharmacokinetic/pharmacodynamic model (MBM). Six clinical CRKP isolates were used for static concentration time kill (SCTK) assays to evaluate the rate and extent of killing by monotherapy, double and triple combinations using PMB, meropenem and fosfomycin. A MBM was developed using the SCTK data in S-ADAPT. The MBM estimated lower maximum killing rate constant of PMB (3.61 h⁻¹) in an isolate with non-functional MgrB and high-level phenotypic resistance. Based upon model discrimination and PMB’s outer membrane disruption, mechanistic synergy was included in 3 isolates which has porin mutations. Mechanistic synergy of PMB was 83-88% with meropenem and 81-98% with fosfomycin. The PMB concentration required to achieve 50% of synergy was 0.48-0.64 mg/L. Simulations with a lower PMB regimen (1mg/kg q12h) and fosfomycin (8g q8h) showed >73% reduction in area under the bacterial load-versus-time curve for four isolates. The triple combination showed 67.7% reduction in non-carbapenamase producing isolate. This study demonstrates that a low dosing regimen of PMB can produce synergistic effects in combination therapy and might be effective in managing infections caused by CRKP, including PMB resistant isolates.

**Author summary:** Antimicrobial resistance is a major concern in treating infectious diseases. CRKP bacterial isolates are resistant to most of the novel antimicrobial agents. One of the primary resistant mechanisms is restricting the permeability of drugs to the site of action. Polymyxin B, an antimicrobial agent, disrupts the bacterial outer membrane, enhancing the permeability of other drugs. When using combination therapy with polymyxin B, selecting drugs with different mechanisms of action is crucial to enhance synergistic effects and improve treatment efficacy. In our study we chose to evaluate the efficacy of meropenem and fosfomycin in double and triple combination therapies. Mechanism based models (MBMs) are the strongest tool to analyse the time course bacterial load data. Our study provides insights into applying available phenotypic and genotypic information to refine and enhance the accuracy of MBMs. Final model simulations revealed that low exposure of polymyxin B below the nephrotoxic threshold was sufficient to produce synergistic effects when combined with fosfomycin and meropenem. Additionally, we found that polymyxin B combination with fosfomycin was more effective compared to meropenem in treating CRKP.

## Introduction

Infections caused by carbapenem-resistant Enterobacterales (CRE) are a significant public health crisis, as the development of new antibiotics is not keeping pace with the rapid rise in antimicrobial resistance (AMR). The global incidence of CRE has doubled over two years, increasing from an average of 4.2% in 2018 to 8.04% between 2020 and 2022 [1]. In the United States, 83% of clinical CRE isolates are carbapenemase-producing, with the most prevalent carbapenamase genes being *bla*_KPC_ (80%), followed by *bla*_NDM_ (15%), *bla*_IMP_ (5%), and *bla*_OXA-48_ (7%). From 2019 to 2021, there was a 1.3-fold decrease in *bla*_KPC_ and a 5-8-fold increase in isolates with other genes. Non-carbapenamase-producing CRE have other β-lactamase genes (ESBL), outer membrane porin disruptions and/or overexpression of genes encoding efflux pumps [2].

A comprehensive meta-analysis has demonstrated that integrating rapid molecular diagnostics (RMDs) to identify pathogens responsible for bloodstream infections with antimicrobial stewardship programs significantly lowers mortality rates, reduces the time to effective therapy, and shortens hospital stays [3]. Recent studies, PRIMERS I and II (Platforms for Rapid Identification of MDR-Gram negative bacteria and Evaluation of Resistance Studies), have evaluated the accuracy of RMD platforms in identifying β-lactamase (*bla*) genotypes, which confer β-lactam resistance, to assist in the appropriate selection of β-lactams [4]. Collectively, these studies underscore the importance of RMDs in the global effort to combat AMR.

The recent approval of novel β-lactamase inhibitors—including avibactam, relebactam, vaborbactam, taniborbactam, and enmetazobactam—has significantly contributed to reducing CRE-related infections. Other inhibitors like ledaborbactam, zidebactam, xeruborbactam, funobactam and nacubactam are nearing approval or in the late stages of development [5]. However, resistance to these β-lactam/β-lactamase inhibitors have already been observed [6]. Key resistance mechanisms include mutations in the *bla*_KPC_ gene, changes in outer membrane permeability, and *bla*_NDM_, which complicate the management of CRE infections [2].

The wide range of resistant mechanisms in CRE makes it challenging to identifying appropriate therapies managing CRE infections. In the absence of novel antimicrobial agents, optimizing the use of existing antibiotics is a crucial strategy to combat AMR [7]. A multi-study analysis of patients infected with carbapenem-resistant *Klebsiella pneumoniae* (CRKP) found that combination therapy (25%–31%) reduced mortality rates by 2-to 2.35-fold compared to monotherapy (50%–73%) [8–10]. Polymyxin B-based combination therapies are recommended by the current polymyxin guidelines as an effective alternative for CRE infections [11]. However, due to the nephrotoxic effects of polymyxins, their use is not highly favored. Recent clinical trials, such as OVERCOME and AIDA, have shown reduced mortality with colistin-based combination therapy (with meropenem) compared to monotherapy for CRE isolates [12,13].

Despite these advancements, clinicians still face challenges in selecting the most appropriate antibiotic combinations. Fosfomycin, a bactericidal drug with a good safety profile and a different mechanism of action (inhibiting the peptidoglycan synthesis at an earlier stage) compared to β-lactams, is a promising option for combination therapy against Gram negative pathogens. Additionally, fosfomycin has shown *in vitro* and *in vivo* synergy with both meropenem and polymyxin B [14].

Given that an antibiotic’s spectrum of activity is dictated by the resistance mechanisms (carbapenemase producing CRKP vs. non-carbapenamase producing CRKP), a systematic and rational approach that considers these resistance mechanisms alongside the susceptibility profile is necessary for effective management of CRE infections.

Static concentration time-kill assays were conducted to evaluate the impact of treatment with polymyxin B, meropenem and fosfomycin as monotherapies, as well as polymyxin B based combinations with either fosfomycin or meropenem, and a triple combination of all three drugs against six different carbapenamase producing and non-carbapenamase producing CRKP clinical isolates over 24 hours. The strains were genomically characterized to elucidate mechanistic insights into their killing activity based on the resistance mechanisms expressed. The objective of this study was to incorporate mechanistic insights gained from phenotypic and genotypic resistance mechanisms to develop a mechanism-based pharmacokinetic/pharmacodynamic (PK/PD) model.

## Results

### Susceptibility and Resistance Gene Profiles for Clinical Isolates

The minimum inhibitory concentrations (MICs) and relevant resistance genes for the six isolates are summarized in **Table 1**. Among the six isolates, four isolates (BRKP61, BRKP67, BRKP76, BRKP28) expressed *bla*_KPC-2_ with outer membrane porin mutations. Additionally, the MgrB protein in BRKP67 and BRKP28 was non-functional, causing resistance to polymyxin B. The non-carbapenemase producer, KP0016-1 had outer membrane porin mutations, while KP0052-1, was the only isolate that expresses *bla*_NDM_ without any porin mutations. The complete gene characterization profiles for each isolate can be found in **Table S2**.

**Table 1.** Antimicrobial resistance genes and MICs for each of the six CRKP isolates.

|  | Resistance Genes |  |  |  |  |  |  |  |
| --- | --- | --- | --- | --- | --- | --- | --- | --- |
| Isolate | Carbapenamase | *mgrB* | *ompK35* | *ompK36* | *ompK37* | PMB MIC | FOF MIC | MEM MIC |
| BRKP28 | *bla*<sub>KPC-2</sub> | NF | NF | NF | Present | >128 R | 128 R | 256 R |
| BRKP61 | *bla*<sub>KPC-2</sub> | - | NF | NF | Present | <0.5 I | 256 R | 128 R |
| BRKP67 | *bla*<sub>KPC-2</sub> | NF | NF | NF | Present | 8 R | 32 R | 64 R |
| BRKP76 | *bla*<sub>KPC-2</sub> | - | NF | NF | NF | <0.5 I | 32 R | 64 R |
| KP0016-1 | - | - | NF | NF | NF | <0.5 I | 64 R | 64 R |
| KP0052-1 | *bla*<sub>NDM-4</sub> | - | Present | Present | Present | < 0.5 I | 64 R | 64 R |

NF- Non-functional. Polymyxin B and meropenem MICs was interpreted using CLSI breakpoints
for *Klebsiella pneumoniae*. Fosfomycin MICs were interpreted using EUCAST breakpoint for
*E.coli*.

### Evaluation of Pharmacodynamic Effect with Mono and Combination Therapy

**Fig 1A** summarizes the percentage of bactericidal activity and extent of bacterial reduction (i.e., AUC_CFU) observed with each polymyxin B-based double and triple combination. A progressive increase in bactericidal activity was noted from polymyxin B-meropenem to polymyxin B-fosfomycin to the triple drug combination. Increasing polymyxin B concentration from 2 to 4 mg/L led to enhanced bactericidal activity across all strains in combination with meropenem (PMB-MEM: 0 to 33%) and fosfomycin (PMB-FOF: 33 to 67%) and the triple combination (PMB-MEM-FOF: 67 to 83%). Overall, the triple combination showed greater reduction in bacterial burden compared to the double combinations, based on the decrease in AUC_CFU. The bacterial reduction was similar between the higher concentration double combination (PMB 4mg/L + FOF 150mg/L) and the lower concentration triple combination (PMB 2mg/L + MEM 40mg/L + FOF 75mg/L), with comparable median AUC_CFU (range: 32.5 - 37.2 log_10_ CFU/mL·h).

**Fig 1.**
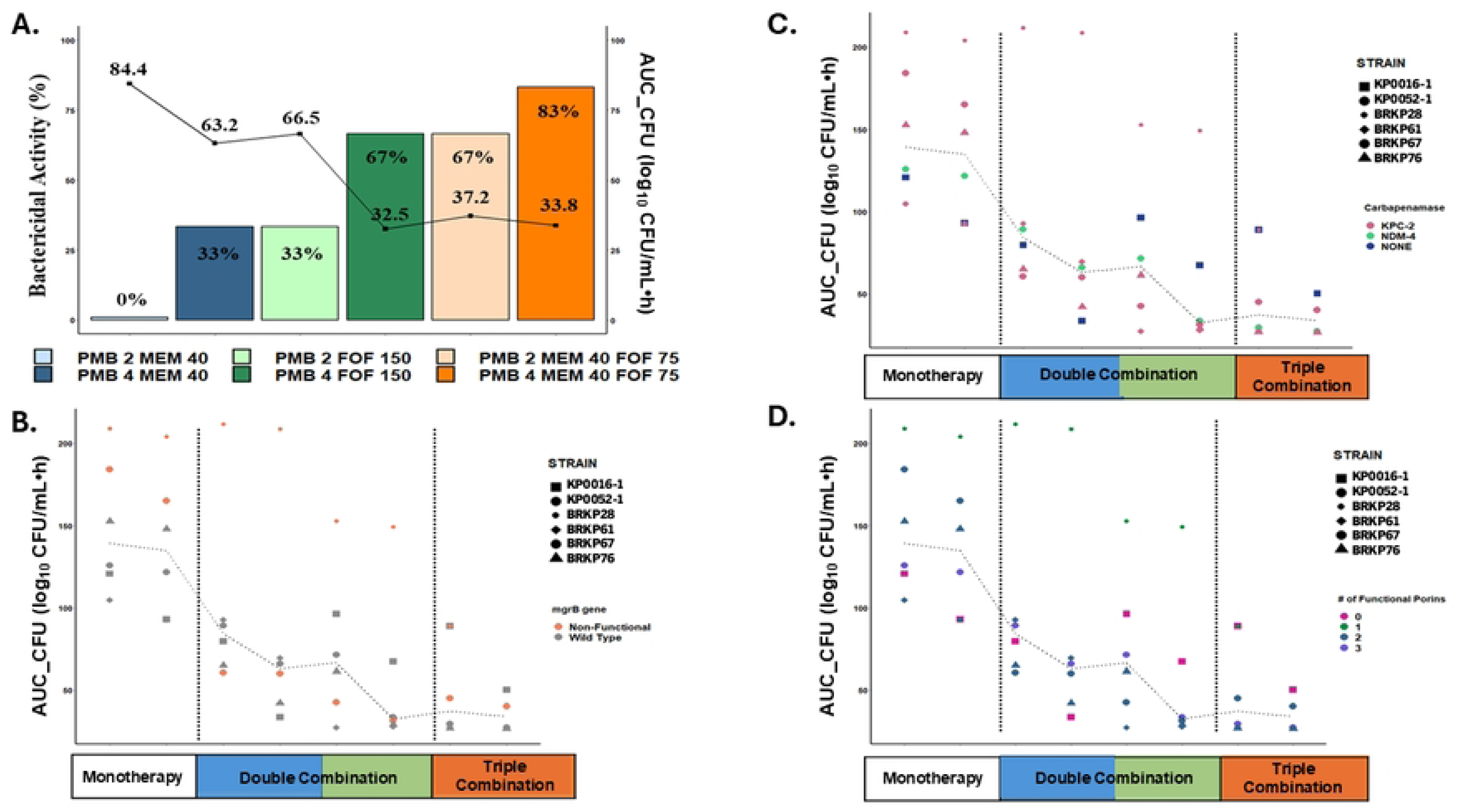
*Panel A* illustrates the bactericidal activity and extent of bacterial reduction (AUC_CFU) for polymyxin B-based double and triple combinations. The bar graph represents the percentage of bactericidal activity, while the line graph shows the median AUC_CFU achieved by each drug regimen. *Panels B-D* shows the corresponding AUC_CFU for each of the six CRKP isolates, categorized by their respective resistance mechanisms: *mgrB* protein (B), carbapenemase production (C) and the number of functional porins (D).

Polymyxin B-based combination therapy was effective against isolates expressing carbapenemases and outer membrane porin mutations (**Fig 1C & D**). Significant pharmacodynamic activity was observed with polymyxin B combination therapy against KP0016-1, even in the absence of functional porins. Against polymyxin B-resistant isolates, double combinations resulted in substantial bacterial reduction against BRKP67 but minimal activity against BRKP28 (**Fig 1B**). The triple combination with higher polymyxin B concentrations (PMB 4mg/L + MEM 40mg/L + FOF 75mg/L), resulted in comparable bacterial reduction against both BRKP67 and BRKP28 (40.1 vs. 50.3 log_10_ CFU/mL·h).

### Mechanism-Based PK/PD Model

The mechanism-based model structure shown in **Fig 2** describes the time-kill data for six isolates, incorporating subpopulation synergy for all of them. When comparing the maximum killing rate constant for polymyxin B resistant isolate BRKP28, which has a non-functional MgrB protein and a high polymyxin B MIC (>128 mg/L), exhibited the lowest maximum killing rate constant for polymyxin B, *Kmax_PMB_*(3.61 h⁻¹) compared to the rest of the isolates (6.61–13.42 h⁻¹). Interestingly, BRKP67 with the *mgrB* mutation and a polymyxin B MIC of 8 mgmg/L had a slightly higher *Kmax_PMB_* (9.07 h⁻¹). Despite both isolates having mutations in MgrB protein, the killing effect of polymyxin B was lower in BRKP28, possibly due to uncharacterized resistance mechanisms contributing to its extremely high MIC (>128 mg/L). The non-carbapenemase-producing KP0016-1 isolate, which has three porin mutations, showed the lowest killing rate constant for both meropenem (1.56 h⁻¹) and fosfomycin (0.67 h⁻¹).

**Fig 2.**
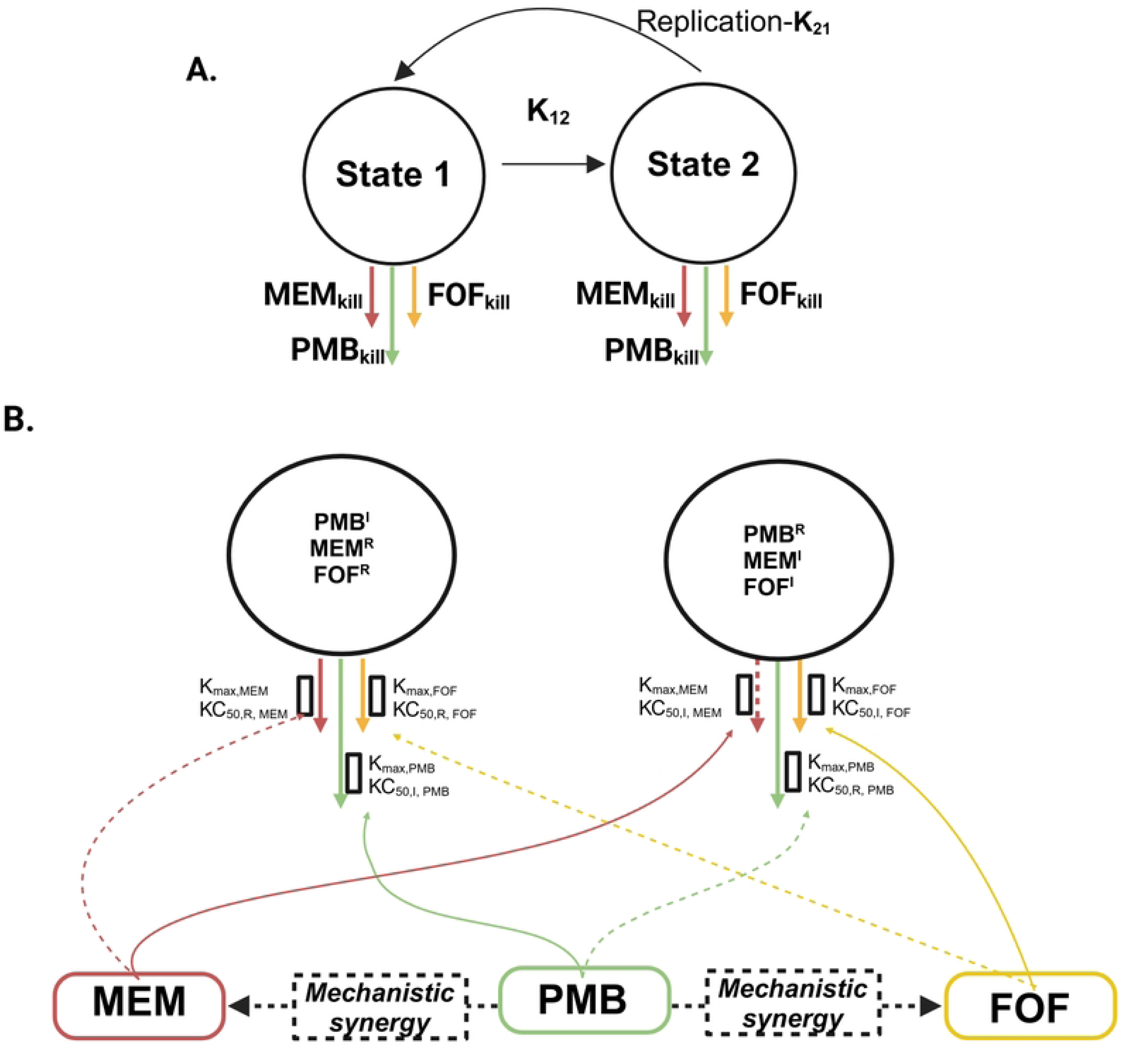
The mechanism-based PK/PD model structure describing the bacterial killing by meropenem (MEM_kill_), polymyxin B (PMB_kill_) and fosfomycin (FOF_kill_) as monotherapy and in combination therapy. (A) A two-state bacterial life cycle model was utilized to describe the bacterial replication and drug effect of bacterial killing is on both state 1 and state 2 of life cycle model. (B) The first subpopulation PMB^I^/MEM^R^/FOF^R^, is intermediately resistant to polymyxin B (PMB) and resistant to meropenem (MEM) and fosfomycin (FOF). The second subpopulation PMB^R^/MEM^I^/FOF^I^, is resistant to polymyxin B (PMB) and intermediately resistant to meropenem (MEM) and fosfomycin (FOF). The maximum killing rate constants (K_max_) and the associated concentrations causing 50% of K_max_ (KC_50_) are explained in **Table 2**. The structure also depicts the shift in meropenem (MEM) and fosfomycin (FOF) KC_50_ resulting from the mechanistic synergy because of polymyxin B (PMB) impacting MEM/FOF – intermediate/resistant subpopulation.

**Table 2.**
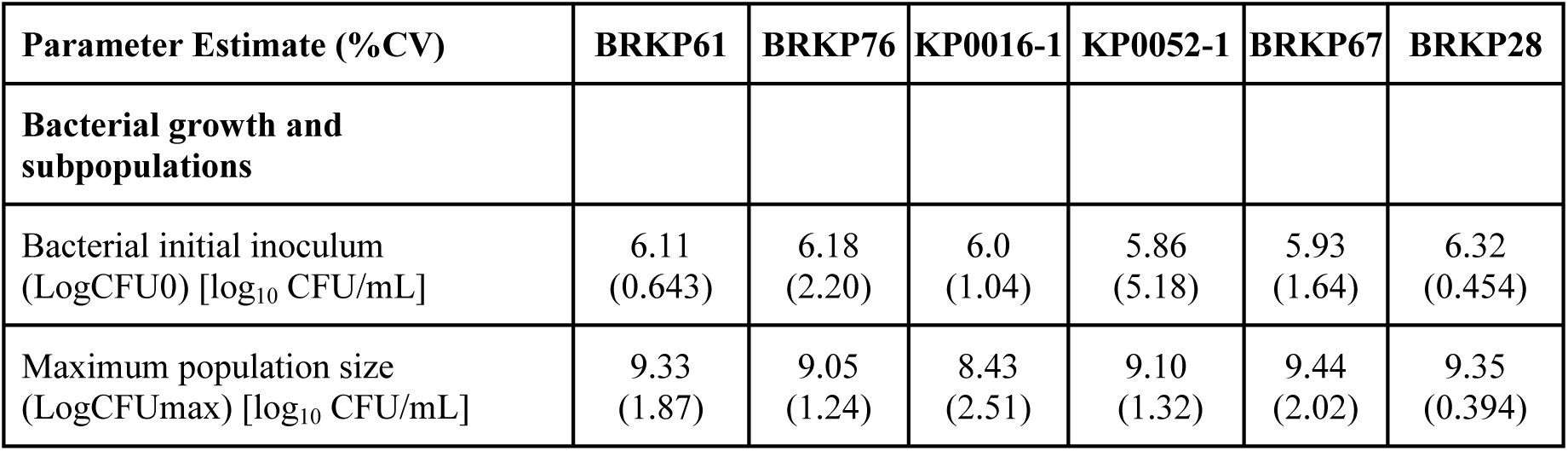
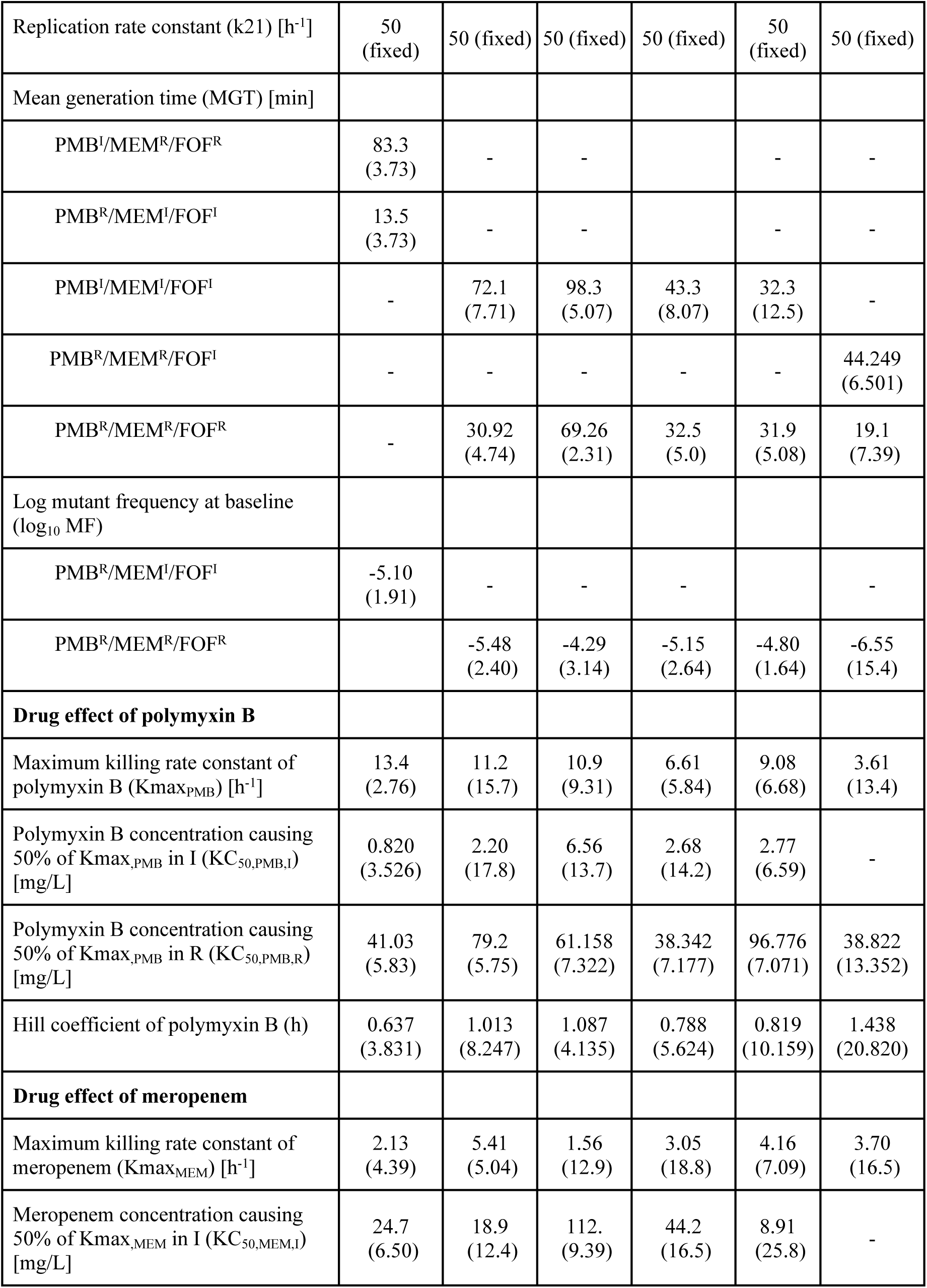
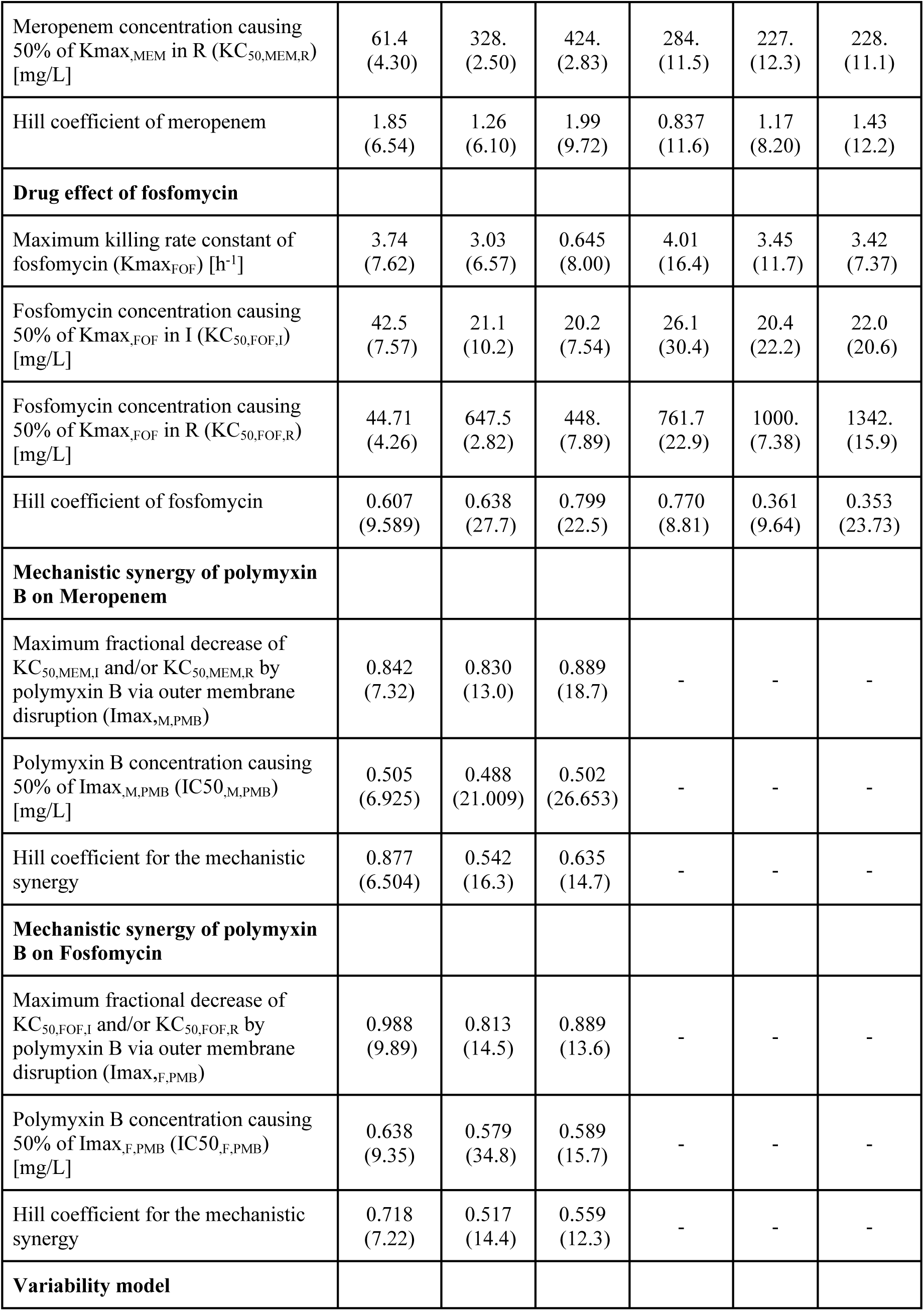
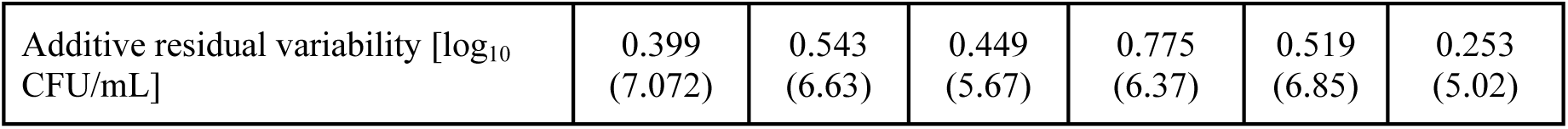
Final model parameter estimates.

Among isolates with a similar meropenem MIC of 64 mg/L, *bla*_NDM-4_ producers had a lower *Kmax* for meropenem (3.05 h⁻¹) compared to *bla*_KPC-2_ producers (4.15–5.44 h⁻¹). Other *bla*_KPC-2_ producers, such as BRKP61 (MIC: 128 mg/L) and BRKP28 (MIC: 256 mg/L), with higher MICs showed lower *Kmax* values for meropenem (2.13 h⁻¹ and 3.69 h⁻¹, respectively).

Through model discrimination, mechanistic synergy was included only for the isolates BRKP61, KP0016-1, and BRKP76, which have mutations in the outer membrane porins. These mutations are assumed to reduce the penetration of hydrophilic drug molecules like meropenem and fosfomycin. Consequently, polymyxin B’s effect on the outer membrane is expected to increase the target site exposure for both drugs. The mechanistic synergy of polymyxin B with meropenem had *Imax,_M,PMB_* values of 0.84, 0.83, and 0.88 for BRKP61, BRKP76 and KP0016-1 respectively. The polymyxin B concentration required to achieve 50% of *Imax,_M,PMB_*ranged from 0.48 to 0.51 mg/L. For fosfomycin, the *Imax,_F,PMB_* values were 0.98, 0.81, and 0.89 for BRKP61, BRKP76, and KP0016-1 respectively, with a polymyxin B concentration of 0.58-0.64 mg/L required to reach 50% of *Imax,_F,PMB_*.

In contrast, for isolates BRKP67 and BRKP28, which express non-functional MgrB protein and have outer membrane porin mutations, including mechanistic synergy did not improve the model. According to the resistance gene profiles and model discrimination, the KP0052-1 isolate, which lacks outer membrane porin mutations, was well described by subpopulation synergy alone. The correlation coefficients for the observed versus model predicted log_10_ CFU/mL were 0.79, 0.73, 0.84, 0.74, 0.73 and 0.91 for BRKP61, BRKP76, KP0016-1, KP0052-1, BRKP67 and BRKP28, respectively (**S2 Fig**). Parameter estimates for each isolate are shown in **Table 2**. The model predicted time course bacterial load data for triple combination (**Fig 3),** double combination (**S3 Fig**) and monotherapy (**S4 Fig**) align well with the observed data.

**Fig 3.**
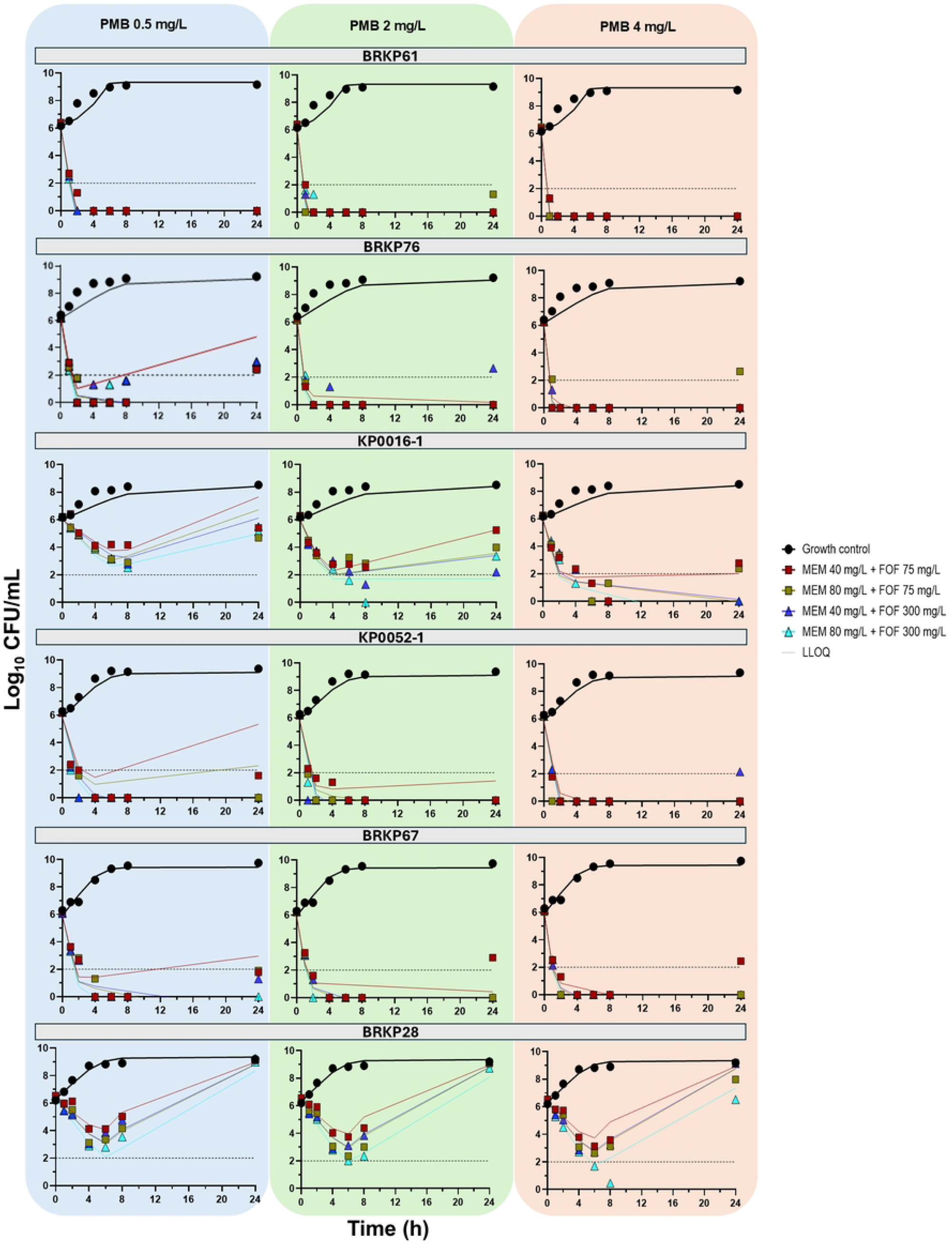
The model predictions for triple combination therapy with polymyxin B 0.5 mg/L, 2 mg/L and 4 mg/L for six isolates are shown. The solid line represents the model predictions, while the symbols indicate the observed data. The horizontal dotted line marks the LLOQ (2 log_10_ CFU/mL).

### Model Predicted Bacterial Load Reduction in Critically ill Patients

Model simulations for critically ill patients using a population PK model showed median polymyxin B exposures of 39.7 mg·h/L and 49.0 mg·h/L for the recommended regimens of LD 2 mg/kg + MD 1.25 mg/kg q12h and LD 2.5 mg/kg + MD 1.5 mg/kg q12h, respectively. The previous population PK model recommended a fixed dosing regimen of LD 150mg + MD 75mg q12h, resulting in a polymyxin B exposure of 39.9 mg·h/L similar to the lower range of recommended weight-based regimen [15]. A slightly lower polymyxin B regimen of 1 mg/kg q12h resulted in an exposure of 23.5 mg·h/L, which is 41-52% lower than the recommended regimens. **Fig 4** presents box plots showing the percentage reduction in AUC_CFU for six isolates treated with double and triple combination therapy, and **S5 Fig** displays the time course log_10_ CFU/mL data for the simulation.

- Green and blue triangle represents isolates with *bla*_KPC-2_ and *bla*_NDM-4_ respectively.
- Orange, purple and brown circle indicate the presence of *ompK-35, ompK-36* and *ompK-37* mutations respectively.
- The brown square represents the insertion of ISKpn13 (1148 bp), an IS5-like element in the *mgrB*, with a polymyxin B MIC of 8 mg/L.
- The red square represents a premature stop codon in the MgrB protein, with a polymyxin B MIC of >128 mg/L.

**Fig 4.**
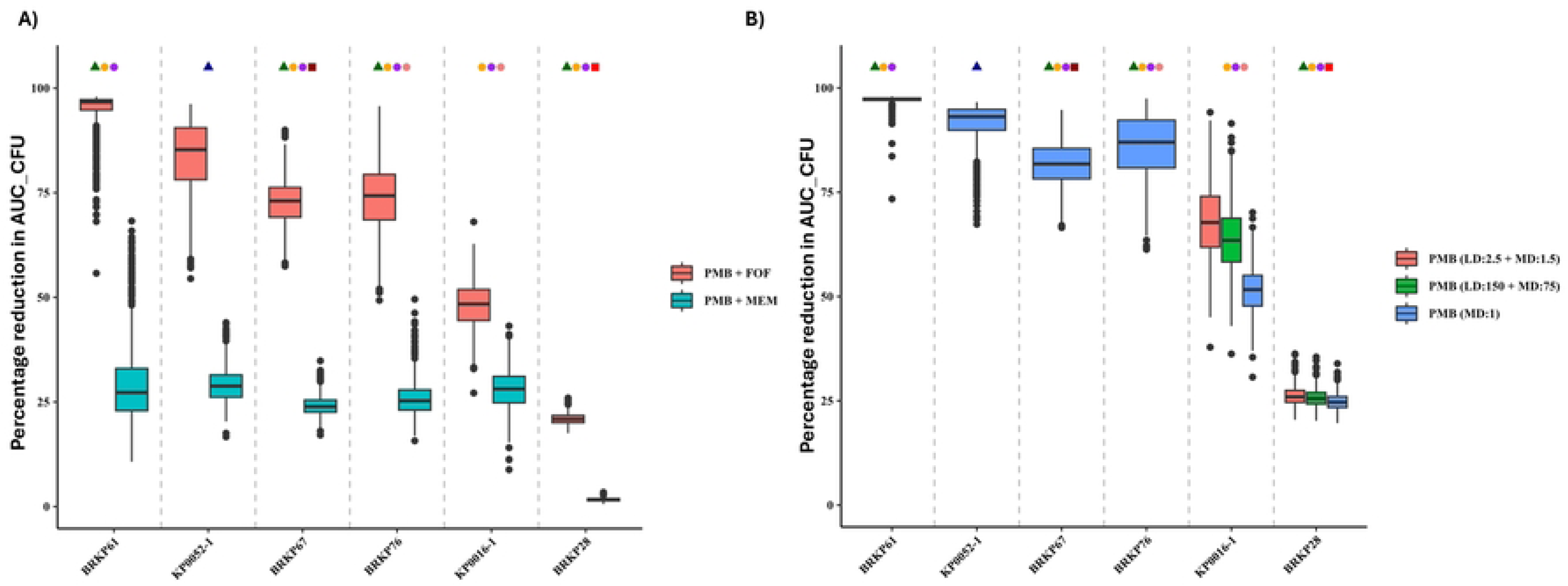
Box plots showing the percentage reduction in AUC_CFU for **A)** double combination therapy with polymyxin B (1 mg/kg q12h) + meropenem (2g q8h) or fosfomycin (8g q8h) **B)** Box plots showing the percentage reduction in AUC_CFU for triple combination therapy with different polymyxin B regimens. Fosfomycin was dosed at 4g q8h for BRKP61 and 8g q8h for all other isolates. Meropenem was dosed at 2g q8h for two isolates KP0016-1 and BRKP28 and 1g q8h for all other isolates.

The double combination of polymyxin B (1mg/kg q12h) with meropenem (2g q8h) showed less than 30% reduction in AUC_CFU for all six isolates. For the BRKP61 isolate, which expresses the carbapenemase enzyme *bla*_KPC-2_ along with two outer membrane porin mutations, the double combination of polymyxin B (1mg/kg q12h) with fosfomycin (8g q8h) resulted in a 96.7% reduction. The triple drug combination was effective (97.3% reduction) when the dose of fosfomycin was lowered (4g q8h), along with lower doses of meropenem (1g q8h) and polymyxin B (1mg/kg q12h). Similarly, the KP0052-1 strain, harboring the *bla*_NDM-4_ without porin mutations, showed an 85.3% reduction with the polymyxin B (1mg/kg q12h) and fosfomycin (8g q8h) combination. The triple combination with same doses of polymyxin B and fosfomycin but a lower dose of meropenem (1g q8h) resulted in 93.1% reduction in AUC_CFU.

The double combination of polymyxin B (1mg/kg q12h) and fosfomycin (8g q8h) showed 74.3% and 73.0% reductions for the BRKP76 (*bla_KPC-2_* and three porin mutations) and BRKP67 (*bla_KPC-2_*, two porin and *mgrB* mutations) isolates respectively. The triple combination with the addition of lower meropenem dose (1g q8h) improved the percentage reduction of AUC_CFU by ∼1.1 fold (BRKP76: 87.0% and BRKP67: 81.8%).

For the non-carbapenemase-producing strain KP0016-1, which also has three porin mutations, the triple drug combination with median polymyxin B exposure (23.5 mg·h/L) resulted in only a 51.7% reduction in AUC_CFU. Increasing the median exposure to 49.0 mg·h/L enhanced the reduction to 67.7%. The BRKP28 strain, which has a premature stop codon in the *mgrB* gene and a high polymyxin B MIC of >128 mg/L, showed only a minimal reduction of 20.9% with polymyxin B and fosfomycin. Furthermore, no significant reduction was observed with the triple drug combination. **Fig 5** illustrates the overall workflow for determining polymyxin B combination therapy based on carbapenamases production and polymyxin B susceptibility.

**Fig 5.**
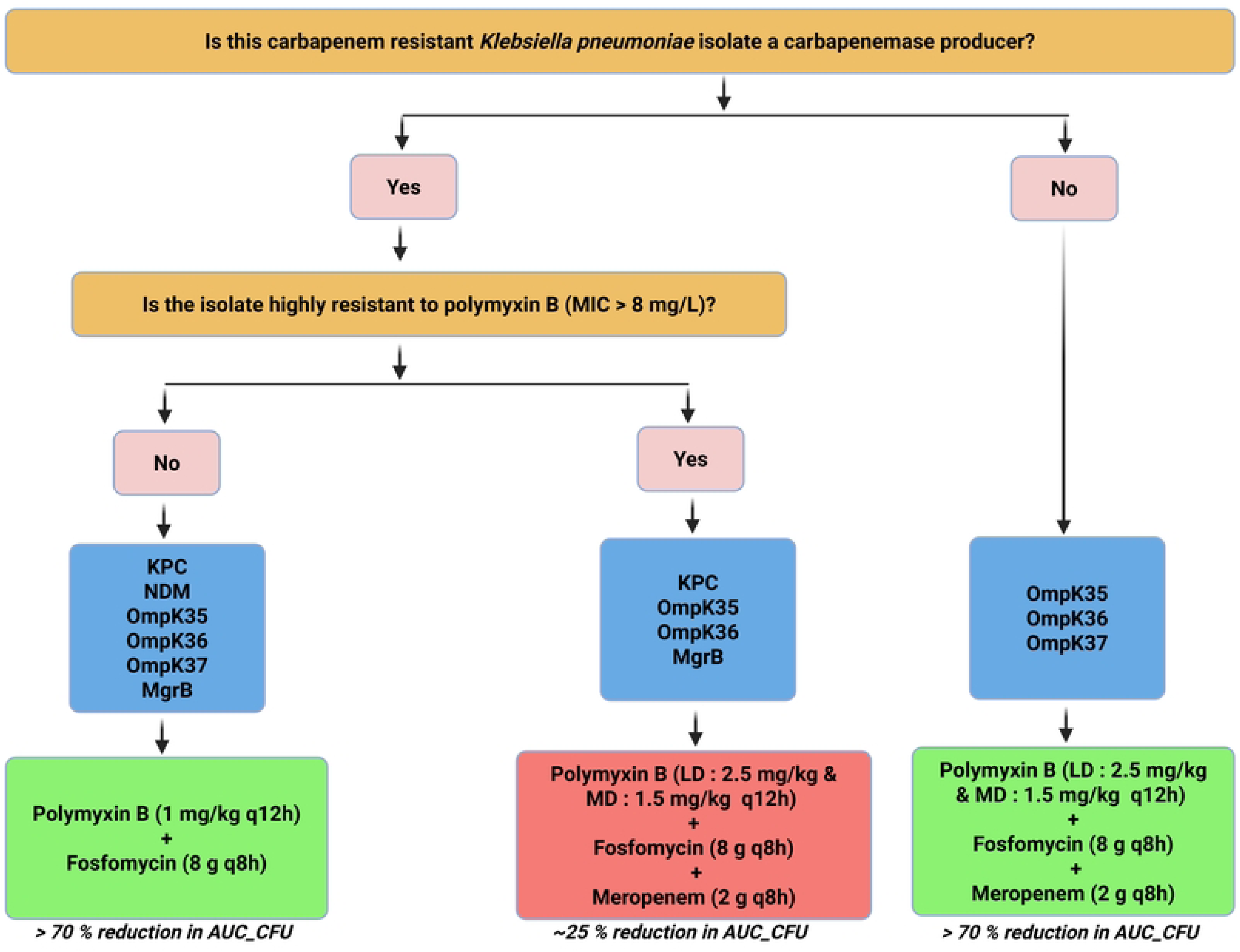
Workflow for selecting polymyxin B based combination therapy. The decision pathway is based on carbapenemases production and polymyxin B susceptibility. Yellow boxes represent the critical questions used to determine the treatment regimens. Pink boxes represent decision points. Blue boxes show the different resistance genes present among the six isolates. Green boxes represent treatment regimens that were successful, achieving >70 % reduction in AUC_CFU. Red box indicate unsuccessful treatment regimens, with only a 25% reduction in AUC_CFU.

## Materials and methods

### Antimicrobials and media

Mueller-Hinton broth (MHB; Becton Dickinson, Franklin Lakes, NJ) supplemented with calcium and magnesium (CAMHB; 25.0 mg/L Ca^2+^, 12.5 mg/L Mg^2+^) and Mueller-Hinton II agar (MHA; Becton, Dickinson, Franklin Lakes, NJ) were used for susceptibility testing and all *in vitro* models. CAMHB was supplemented with 25 mg/L glucose-6-phosphate (lot number 343234, Acros Organics). Stock solutions of polymyxin B (lot number WXBB5309V; Sigma Aldrich, St. Louis, MO), meropenem (lot number LC24337; AK Scientific, Union City, CA) and fosfomycin (lot number: K001, Nabriva Therapeutics US, Inc., Fort Washington, PA) were freshly prepared in sterile water and saline prior to each experiment. All drug solutions were filter sterilised with a 0.22 µm filter (Fisher Scientific, Pittsburgh, PA).

### Bacterial isolates and antibiotic susceptibility testing

Four clinical carbapenem-resistant *Klebsiella pneumoniae* isolates used in this study (BRKP28, BRKP61, BRKP67, BRKP76) were obtained from different patients from Instituto Dante Pazzanese de Cardiologia (Sao Paulo, Brazil) and two isolates (KP0016-1, and KP0052-1) were obtained from Siriraj Hospital (Bangkok, Thailand) [16]. MICs for polymyxin B and meropenem were determined in triplicate using the broth microdilution method, whereas fosfomycin susceptibility testing was performed using the agar dilution method, both according to CLSI guidelines [17]. MICs for polymyxin B and meropenem were interpreted using CLSI breakpoints for *K. pneumoniae.* Due to the lack of CLSI breakpoints for fosfomycin against *K. pneumoniae*, we used breakpoints provided by EUCAST for fosfomycin IV against *Escherichia coli*.

### Genomic characterization

Polymerase chain reaction (PCR) was performed using previously described primer sets for β-lactamase Ambler classes A (*bla*_GES_ and *bla*_KPC_), B (*bla*_NDM-1_, *bla*_VIM_, and *bla*_IMP_), and D (*bla*_OXA-48_ and *bla*_OXA-40_) [18]. Primers used in amplification experiments to identify *mgrB* [19] and *ompK* genes [20] (*ompK35*, *ompK36* & *ompK37*) are listed in **Table S1**. Isolates were also characterized for the virulence genes and the primers used for the virulence genes are mentioned in **Table S1**. Genomic DNA was extracted from bacterial isolates using the Purelink™ Genomic DNA Mini kit (Invitrogen, Carlsbad, CA). PCR reactions were performed using Q5 Hi-Fidelity Taq DNA Polymerase (New England Biolabs, Ipswich, MA). Reactions were carried out in an Eppendorf Mastercycler® (Eppendorf, Hamburg, Germany), and PCR products were analysed by direct sequencing (GENEWIZ, Research Triangle Park, NC). Nucleotide and deduced protein sequences were analysed using BLAST (http://blast.ncbi.nlm.nih.gov/).

### Static concentration time-kill (SCTK) Assays

SCTK assays were performed over 24 hours to evaluate the rate and extent of killing by monotherapy, double and triple combinations using polymyxin B, meropenem and fosfomycin. PK/PD analysis were performed using six isolates at a starting inoculum of ∼10^6^ CFU/mL (CFU_0_), as previously described [21]. The antibiotic concentrations selected represent a combination of clinically achievable (polymyxin B: 0.5, 1, 2, 4, 8 mg/L; meropenem: 10, 20, 40 mg/L; Fosfomycin: 75, 150, 300, 500 mg/L) and supra-therapeutic (polymyxin B: 16 and 64 mg/L; meropenem: 60 and 120 mg/L) free-drug concentrations (i.e., unbound plasma) in order to evaluate the concentration-response relationship against each isolate [15,22,23]. Serial samples were obtained at 0, 1, 2, 4, 6, 8, and 24 h for bacterial quantification. The lower limit of quantification (LLOQ) was 2 log_10_ CFU/mL.

### Pharmacodynamic analysis

The pharmacodynamic effect was quantified as the change in log_10_ CFU/mL at 24 h (CFU*_24_*) compared to baseline (CFU_0_) (i.e., 24h log_10_ CFU/mL reduction). Bactericidal activity was defined as a ≥3 log_10_ reduction at 24h compared to the initial inoculum. To further assess bacterial killing in the polymyxin B-based combination therapy, the area under the bacterial load-versus-time curve (AUC___CFU) from 0 to 24 hours was calculated for both the double and triple drug combinations at static concentrations.

### Mechanism based PK/PD model development

Mechanism-based model was developed using the SCTK time course data describing the change in bacterial burden over time in response to antibiotic treatment against the six bacterial isolates. A life cycle model was used to describe the bacterial growth and replication for each bacterial isolate [24]. The model included the transition of bacterial cells from vegetative state, preparing for replication (state 1), to the replication state, immediately prior to replication step (state 2).

A mixture model with pre-existing subpopulations with differing susceptibilities to polymyxin B, meropenem and fosfomycin were considered to account for heterogeneity in the bacterial inoculum. Models with two, three and four bacterial subpopulations were explored [25]. The final model included two subpopulations with differing susceptibilities to polymyxin, meropenem and fosfomycin for each isolate (1).

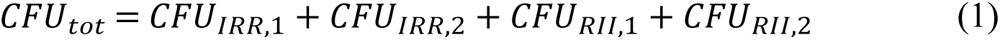

where *CFU_tot_* is the total viable bacterial concentration, *CFU_IRR_* is the polymyxin B-intermediate, meropenem-resistant and fosfomycin-resistant subpopulation. *CFU_RII_* is the polymyxin B-resistant, meropenem-intermediate and fosfomycin-intermediate subpopulation. Each of the bacterial subpopulations, *CFU_IRR,1_* and *CFU_IRR,2_* is in vegetative (state 1) and replicative state (state 2). The differential equation for the two states describing the bacteria in state 1 and 2 for subpopulation *CFU_IRR_* with killing by polymyxin B (PMB), meropenem (MEM), and fosfomycin (FOF) are shown in equation (2) and (3).

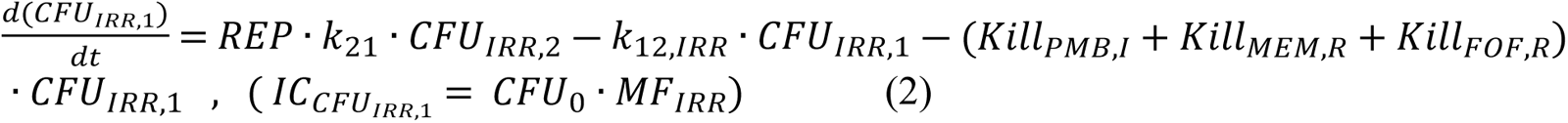

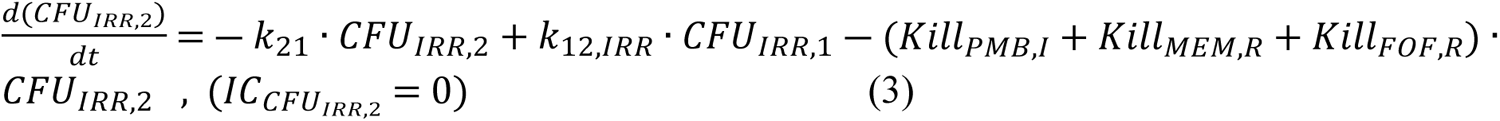

where REP is the replication factor defined as 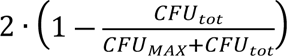, *CFU_MAX_* is the maximum bacterial population size and 2 represents the doubling of bacteria during replication. The inverse of the mean replication time from state 2 to state 1, *k_21_* was fixed to 50 h^-1^. *k_12,IRR_*, the inverse of mean replication time from state 1 to state 2 was estimated for each subpopulation. *Kill_PMB,I_*, *Kill_MEM,R_* and *Kill_FOF,R_* are the killing rate by polymyxin B, meropenem and fosfomycin respectively for *CFU_IRR_*subpopulation. The killing activity of each drug is described by the Hill equation as shown in equation (4)

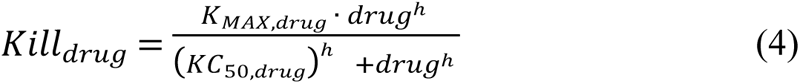

where drug can be concentration of polymyxin B, meropenem or fosfomycin, *K_MAX,drug_* is the maximum killing rate constant of drug, *KC_50,drug_* is the drug concentration causing 50% of *K_MAX,drug_*, ℎ is the Hill coefficient. We considered subpopulation synergy (i.e., polymyxin B killing bacteria resistant to meropenem/Fosfomycin and vice versa) in this model. We estimated the total initial inoculum (log_10_CFU_0_) and the log transformed mutation frequency for the subpopulations. Initial conditions were implemented as described previously [25].

The interaction between polymyxin B and meropenem and/or fosfomycin was modeled using a Hill function to describe the mechanistic synergy based on polymyxin B’s effect on the outer membrane of the Gram-negative bacteria. This synergy increases the target site concentration of meropenem and fosfomycin, thereby enhancing the sensitivity of the intermediate and resistant subpopulations to meropenem and fosfomycin (*KC_50,MEM,I_*, *KC_50,FOF,I_, KC_50,MEM,R_*, *KC_50,FOF,R_*). Equations (5) describes the mechanistic synergy by polymyxin B and equation (6) and (7) describes the impact of mechanistic synergy on meropenem intermediate and fosfomycin resistant subpopulation respectively.

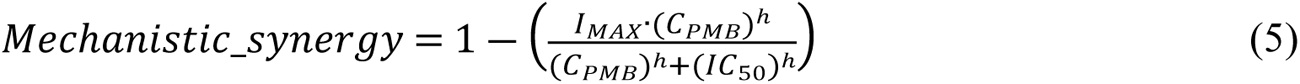

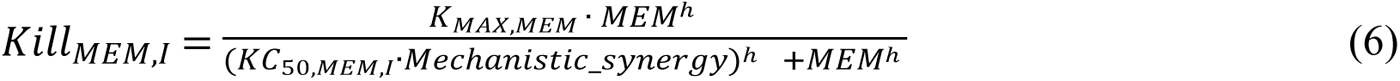

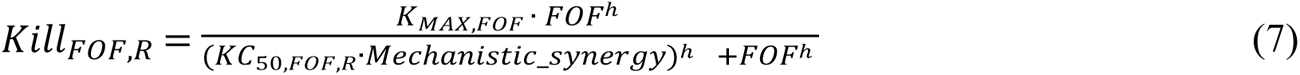

Where *I_MAX_*is the maximum fractional decrease in *KC_50,MEM,I_* and *KC_50,FOF,R_*by polymyxin B causing disruption of the bacterial outer membrane, *IC_50_*is the polymyxin B concentration causing 50% of *I_MAX_*, *h* is the hill coefficient.

The residual unexplained variability for log_10_ transformed bacterial load data was explained by additive error model. Observations below the LLOQ were fit using the Beal M3 [26]. Parameter estimation was performed using the importance sampling algorithm (pmethod=4) in S-ADAPT (version 1.57) facilitated by SADAPT-TRAN [27]. Models were evaluated based on model fits, diagnostic plots, precision of parameter estimates, and objective function value (OFV).

### Monte Carlo simulations to predict drug response in clinical practice

Monte Carlo simulations were performed for 1,000 adult patients using mrgsolve R package. Previously published population PK models for polymyxin B [15], meropenem [22] and fosfomycin [23] in critically ill patients were used to simulate the unbound plasma concentrations taking inter individual variability into consideration **(S1 Fig)**. The unbound plasma concentrations were determined using published fraction unbound (*fu)* for polymyxin B=0.42 and meropenem= 0.98, while plasma protein binding for fosfomycin was negligible. These simulated unbound drug concentrations were linked to the developed MBM to predict impact on bacterial killing in critically ill patients. Simulations were performed using demographics within the reported range in the population PK models for polymyxin B, meropenem and fosfomycin [bodyweight of 70 kg, serum albumin of 2.8 g/dl and creatinine clearance of 90 ml/min].

The efficacy of a lower polymyxin B dosing regimen (1 mg/kg every 12 h as 1 h infusion) was evaluated in both the double and triple combination regimens. In cases where the lower dosing regimen did not result in a significant reduction in AUC_CFU, simulations were performed using the recommended weight-based polymyxin B regimen (2.5 mg/kg loading dose [LD] followed by 1.5 mg/kg every12 h maintenance dose [MD] as 1 h infusion) and a fixed regimen (150 mg LD and 75 mg every 12 h MD as 1 h infusion). Additionally, meropenem (1 g or 2 g every 8 h as 3 h infusion) and fosfomycin (4 g or 8 g every 8 h as 3 h infusion) regimens were simulated. The pharmacodynamic activity of double and triple combination therapy was evaluated as a percentage reduction in AUC_CFU with the treatment of interest compared to no treatment.

## Discussion

The rising incidence of both carbapenemase producing and non-carbapenemase producing CRKP infections, coupled with a lack of effective treatment options, necessitates detailed *in vitro* evaluation of potential treatment regimens using approved drugs. Although newer β-lactams/β-lactamase inhibitor combinations are the recommended treatment for carbapenemase producing CRKP, there is increasing evidence of resistance due to overexpressed efflux pumps, porin loss, or mutations in carbapenemases [28–31]. Several studies have investigated the efficacy of polymyxin B-based combinations against NDM-producing *K. pneumoniae,* highlighting polymyxin’s role as a last-line agent against these difficult-to-treat pathogens [32–34]. Observational clinical data supports the use of meropenem containing combinations due to their lower mortality rates [35]. Another small prospective study found that combination therapy with intravenous fosfomycin reduced all-cause hospital mortality in critically ill patients with CRKP infections [36]. Meropenem and fosfomycin were selected for their different mechanisms of action: meropenem inhibits peptidoglycan synthesis by binding to penicillin-binding proteins, while fosfomycin prevents the transpeptidation of peptidoglycan. These two drugs are hydrophilic in nature, and mutations in outer membrane porins can hinder their ability to penetrate and achieve adequate exposure at the site of action. Polymyxin B’s detergent like activity on the outer membrane enhances the target site concentration of meropenem and fosfomycin [16,32]. Recent Infectious Diseases Society of America (IDSA) guidance does not recommend polymyxin-based combination regimens for CRE infections due to the increased toxicity associated with this narrow therapeutic index drug [2]. Despite this, various clinical trials have shown evidence of reduced mortality in CRE due to polymyxin-based combination therapy [12,13].

To evaluate the synergistic effects of polymyxin B with meropenem and/or fosfomycin, we selected six different isolates with varying phenotypic and genotypic resistance mechanisms. These included strains with carbapenemases like *bla*_KPC-2_ or *bla*_NDM-4,_ with or without porin mutations, which confer resistance to meropenem. Fosfomycin permeates well through *E. coli ompF,* which is a homologue of *K. pneumoniae ompK35* [37]. Five out of six isolates had *ompK35* mutations, which can limit fosfomycin penetration. We also included two isolates with mutations in MgrB protein, one of which exhibited extreme phenotypic resistance to polymyxin B (>128 mg/L).

This information about phenotypic and genotypic resistance mechanisms provided significant insights into the mechanistic synergy in MBM across the six isolates. Based on the presence of porin mutations and polymyxin B susceptibility, mechanistic synergy was applicable to only three isolates. Our model estimated 83-88% mechanistic synergy for polymyxin B with meropenem and 81-98% with fosfomycin. Although there was over 80% synergy of polymyxin B with meropenem compared to monotherapy, it was not significant enough to reduce AUC_CFU. In contrast, the 81-98% synergy of polymyxin B with fosfomycin led to a significant reduction in AUC_CFU in four isolates. In this study, polymyxin B-based combination therapy was effective against all isolates except BRKP28 which showed only minimal activity due to high MIC values due to uncharacterized resistance mechanisms, possibly due to positively charged lipopolysaccharide (LPS) which could interfere with polymyxin B’s interaction to LPS.

The re-emergence of polymyxin B is not without challenges, given the increasing reports of resistance. The absence of *mgrB* disrupts negative feedback to the PhoQ/PhoP signalling circuit, adding a cationic product (4-amino-4-deoxy-L-arabinose) to lipid A in the lipopolysaccharide target of polymyxin B, resulting in resistance [19]. A recent study found that the triple combination of polymyxin B, meropenem and fosfomycin significantly restricts the selective amplification of polymyxin B resistant subpopulations in *Acinetobacter baumannii* [38]. Similarly, our study supports the use of polymyxin B-based combinations against polymyxin-resistant isolates like BRKP67, suggesting that the polymyxin B MIC (i.e., phenotypic susceptibility) may not be predictive of the pharmacodynamic response. MacNair et al. found that *Enterobacteriaceae* strains harbouring *mcr-1* did not impede colistin’s ability to disrupt the outer membrane, indicating that acquired resistance to polymyxins does not necessarily hinder its synergistic potential in combination [39]. Sharma et al. revealed that polymyxin B led to extensive morphological changes in all isolates except BRKP28, which showed minimal changes.[16]

Clinicians are concerned about higher exposure to polymyxin B leading to nephrotoxicity. Tailored polymyxin B-based combination therapy should be considered to ensure adequate drug exposure while minimizing the risk for nephrotoxicity [33]. A retrospective study in adult patients found that the probability of polymyxin B-associated acute kidney injury is 50% for trough concentrations ≥ 3.13 mg/L [40]. Our simulations with the lowest polymyxin B regimen of 1 mg/kg every 12h maintained trough concentrations below the nephrotoxic threshold. Additionally, our model simulations demonstrated that the lowest polymyxin B exposure is sufficient to provide synergistic effects in combination therapy. Given the minimal safety concerns with fosfomycin, the maximum recommended dose (8g every 8h) should be used to benefit from synergistic effects. In some cases where triple combination therapy is needed, addition of low dose meropenem (1g every 8h) or the recommended meropenem regimen (2g every 8h) is necessary to achieve bacterial reduction. Trough concentrations of these regimens were below the nephrotoxic (44.5 mg/L) and neurotoxic (64.5 mg/L) thresholds [41] (**S1 Fig**).

Although our study evaluates the pharmacodynamic activity in a limited number of isolates expressing a range of resistance mechanisms, our findings align with previous systematic reviews addressing therapeutic approaches for multidrug-resistant Gram-negative bacteria [42,43]. The role of novel β-lactamase inhibitors and polymyxin B-based combination therapy should be further explored against a larger panel of CRKP isolates with various genotypes (i.e., non-carbapenemase vs. carbapenemase-producing) *in vitro* and *in vivo* to determine their potential for clinical use. Furthermore, the mechanism-based model can be improved with additional information about multi-omics data, which will help us understand the impact of each resistance mechanism on the pharmacodynamic activity of each drug during monotherapy and in combination.

### Conclusion

While the need for appropriate antimicrobial treatment is widely acknowledged, high-level evidence to guide antibiotic selection for extensively drug resistant CRE isolates is lacking. This study shows that a low dose polymyxin B regimen can produce synergistic effects in combination with the right antibiotic(s). Such combination therapy can be effective in managing infections caused by both carbapenamase producing and non-carbapenamase producing CRKP, including polymyxin B resistant isolates. The MBM informed polymyxin B dosing ensures safety while maximizing its potential to achieve synergistic effects.

Polymyxin B shows greater synergy with fosfomycin compared to meropenem. Polymyxin B-based triple combination therapy is beneficial for non-carbapenamase producing isolates resistant to newer antibiotics. Understanding resistance mechanisms can inform the new treatment regimens that integrate both newer and old antibiotics to achieve adequate exposure at the infection site.

Future research should explore the benefits of genotype-phenotype associations with quantitative data, particularly pharmacodynamic activity in the presence or absence of a specific resistance genes or mechanisms. Given the limited clinical data, a systematic, rational approach is recommended to generate robust nonclinical PK/PD data to ensure the longevity of our antibiotic arsenal.

## References

1. Trusts PC. Antibiotics Currently in Global Clinical Development. https://www.pewtrusts.org/en/research-and-analysis/data-visualizations/2014/antibiotics-currently-in-clinical-development1. Wise MG, Karlowsky JA, Mohamed N, Hermsen ED, Kamat S, Townsend A, et al. Global trends in carbapenem- and difficult-to-treat-resistance among World Health Organization priority bacterial pathogens: ATLAS surveillance program 2018–2022. Journal of Global Antimicrobial Resistance. 2024 Jun 1;37:168–75.

2. Tamma PD, Heil EL, Justo JA, Mathers AJ, Satlin MJ, Bonomo RA. Infectious Diseases Society of America 2024 Guidance on the Treatment of Antimicrobial-Resistant Gram-Negative Infections. Clinical Infectious Diseases. 2024 Aug 7;ciae403.

3. Timbrook TT, Morton JB, McConeghy KW, Caffrey AR, Mylonakis E, LaPlante KL. The Effect of Molecular Rapid Diagnostic Testing on Clinical Outcomes in Bloodstream Infections: A Systematic Review and Meta-analysis. Clin Infect Dis. 2017 Jan 1;64(1):15– 23.

4. Evans SR, Hujer AM, Jiang H, Hujer KM, Hall T, Marzan C, et al. Rapid Molecular Diagnostics, Antibiotic Treatment Decisions, and Developing Approaches to Inform Empiric Therapy: PRIMERS I and II. Clin Infect Dis. 2016 Jan 15;62(2):181–9.

5. Koenig C, Kuti JL. Evolving resistance landscape in gram-negative pathogens: An update on β-lactam and β-lactam-inhibitor treatment combinations for carbapenem-resistant organisms. Pharmacotherapy. 2024 Aug;44(8):658–74.

6. Lin CK, Page A, Lohsen S, Haider AA, Waggoner J, Smith G, et al. Rates of resistance and heteroresistance to newer β-lactam/β-lactamase inhibitors for carbapenem-resistant Enterobacterales. JAC-Antimicrobial Resistance. 2024 Apr 1;6(2):dlae048.

7. NIAID’s Antibiotic Resistance Research Framework: Current Status and Future Directions 2019.

8. Falagas ME, Lourida P, Poulikakos P, Rafailidis PI, Tansarli GS. Antibiotic treatment of infections due to carbapenem-resistant Enterobacteriaceae: systematic evaluation of the available evidence. Antimicrob Agents Chemother. 2014;58(2):654–63.

9. Sheu CC, Chang YT, Lin SY, Chen YH, Hsueh PR. Infections Caused by Carbapenem-Resistant Enterobacteriaceae: An Update on Therapeutic Options. Front Microbiol. 2019;10:80.

10. Qureshi ZA, Paterson DL, Potoski BA, Kilayko MC, Sandovsky G, Sordillo E, et al. Treatment outcome of bacteremia due to KPC-producing Klebsiella pneumoniae: superiority of combination antimicrobial regimens. Antimicrob Agents Chemother. 2012 Apr;56(4):2108–13.

11. Tsuji BT, Pogue JM, Zavascki AP, Paul M, Daikos GL, Forrest A, et al. International Consensus Guidelines for the Optimal Use of the Polymyxins: Endorsed by the American College of Clinical Pharmacy (ACCP), European Society of Clinical Microbiology and Infectious Diseases (ESCMID), Infectious Diseases Society of America (IDSA), International Society for Anti-infective Pharmacology (ISAP), Society of Critical Care Medicine (SCCM), and Society of Infectious Diseases Pharmacists (SIDP). Pharmacotherapy. 2019 Jan;39(1):10–39.

12. Kaye KS, Marchaim D, Thamlikitkul V, Carmeli Y, Chiu CH, Daikos G, et al. Colistin Monotherapy versus Combination Therapy for Carbapenem-Resistant Organisms. NEJM Evidence. 2022 Dec 27;2(1):EVIDoa2200131.

13. Colistin alone versus colistin plus meropenem for treatment of severe infections caused by carbapenem-resistant Gram-negative bacteria: an open-label, randomised controlled trial - PubMed [Internet]. [cited 2024 Nov 5]. Available from: https://pubmed.ncbi.nlm.nih.gov/29456043/

14. Ribeiro AC da S, Chikhani YC dos SA, Valiatti TB, Valêncio A, Kurihara MNL, Santos FF, et al. In Vitro and In Vivo Synergism of Fosfomycin in Combination with Meropenem or Polymyxin B against KPC-2-Producing Klebsiella pneumoniae Clinical Isolates. Antibiotics. 2023 Jan 23;12(2):237.

15. Hanafin PO, Kwa A, Zavascki AP, Sandri AM, Scheetz MH, Kubin CJ, et al. A population pharmacokinetic model of polymyxin B based on prospective clinical data to inform dosing in hospitalized patients. Clin Microbiol Infect. 2023 Sep;29(9):1174–81.

16. Sharma R, Patel S, Abboud C, Diep J, Ly NS, Pogue JM, et al. Polymyxin B in combination with meropenem against carbapenemase-producing Klebsiella pneumoniae: pharmacodynamics and morphological changes. Int J Antimicrob Agents. 2017 Feb;49(2):224–32.

17. M100 Ed34 | Performance Standards for Antimicrobial Susceptibility Testing, 34th Edition [Internet]. Clinical & Laboratory Standards Institute. [cited 2024 Aug 14]. Available from: https://clsi.org/standards/products/microbiology/documents/m100/

18. Monteiro J, Widen RH, Pignatari ACC, Kubasek C, Silbert S. Rapid detection of carbapenemase genes by multiplex real-time PCR. J Antimicrob Chemother. 2012 Apr;67(4):906–9.

19. Poirel L, Jayol A, Bontron S, Villegas MV, Ozdamar M, Türkoglu S, et al. The mgrB gene as a key target for acquired resistance to colistin in Klebsiella pneumoniae. J Antimicrob Chemother. 2015 Jan;70(1):75–80.

20. Kaczmarek FM, Dib-Hajj F, Shang W, Gootz TD. High-level carbapenem resistance in a Klebsiella pneumoniae clinical isolate is due to the combination of bla(ACT-1) beta-lactamase production, porin OmpK35/36 insertional inactivation, and down-regulation of the phosphate transport porin phoe. Antimicrob Agents Chemother. 2006 Oct;50(10):3396– 406.

21. Diep JK, Jacobs DM, Sharma R, Covelli J, Bowers DR, Russo TA, et al. Polymyxin B in Combination with Rifampin and Meropenem against Polymyxin B-Resistant KPC-Producing Klebsiella pneumoniae. Antimicrob Agents Chemother. 2017 Feb;61(2):e02121–16.

22. Ehmann L, Zoller M, Minichmayr IK, Scharf C, Huisinga W, Zander J, et al. Development of a dosing algorithm for meropenem in critically ill patients based on a population pharmacokinetic/pharmacodynamic analysis. Int J Antimicrob Agents. 2019 Sep;54(3):309–17.

23. Wangchinda W, Pogue JM, Thamlikitkul V, Leelawattanachai P, Koomanachai P, Pai MP. Population pharmacokinetic/pharmacodynamic target attainment analysis of IV fosfomycin for the treatment of MDR Gram-negative bacterial infections. J Antimicrob Chemother. 2024 Jun 3;79(6):1372–9.

24. Luterbach CL, Qiu H, Hanafin PO, Sharma R, Piscitelli J, Lin FC, et al. A Systems-Based Analysis of Mono- and Combination Therapy for Carbapenem-Resistant Klebsiella pneumoniae Bloodstream Infections. Antimicrobial Agents and Chemotherapy. 2022 Sep 20;66(10):e00591.

25. Landersdorfer CB, Ly NS, Xu H, Tsuji BT, Bulitta JB. Quantifying subpopulation synergy for antibiotic combinations via mechanism-based modeling and a sequential dosing design. Antimicrob Agents Chemother. 2013 May;57(5):2343–51.

26. Beal SL. Ways to fit a PK model with some data below the quantification limit. J Pharmacokinet Pharmacodyn. 2001 Oct;28(5):481–504.

27. Bulitta JB, Bingölbali A, Shin BS, Landersdorfer CB. Development of a New Pre- and Post-Processing Tool (SADAPT-TRAN) for Nonlinear Mixed-Effects Modeling in S-ADAPT. AAPS J. 2011 Mar 3;13(2):201–11.

28. Wilhelm CM, Antochevis LC, Magagnin CM, Arns B, Vieceli T, Pereira DC, et al. Susceptibility evaluation of novel beta-lactam/beta-lactamase inhibitor combinations against carbapenem-resistant Klebsiella pneumoniae from bloodstream infections in hospitalized patients in Brazil. J Glob Antimicrob Resist. 2024 Sep;38:247–51.

29. Papp-Wallace KM, Mack AR, Taracila MA, Bonomo RA. Resistance to Novel β-Lactam-β-Lactamase Inhibitor Combinations: The “Price of Progress.” Infect Dis Clin North Am. 2020 Dec;34(4):773–819.

30. Sun D, Rubio-Aparicio D, Nelson K, Dudley MN, Lomovskaya O. Meropenem-Vaborbactam Resistance Selection, Resistance Prevention, and Molecular Mechanisms in Mutants of KPC-Producing Klebsiella pneumoniae. Antimicrob Agents Chemother. 2017 Dec;61(12):e01694–17.

31. Humphries RM, Yang S, Hemarajata P, Ward KW, Hindler JA, Miller SA, et al. First Report of Ceftazidime-Avibactam Resistance in a KPC-3-Expressing Klebsiella pneumoniae Isolate. Antimicrob Agents Chemother. 2015 Oct;59(10):6605–7.

32. Abdul Rahim N, Cheah SE, Johnson MD, Yu H, Sidjabat HE, Boyce J, et al. Synergistic killing of NDM-producing MDR Klebsiella pneumoniae by two “old” antibiotics-polymyxin B and chloramphenicol. J Antimicrob Chemother. 2015 Sep;70(9):2589–97.

33. Garcia E, Diep JK, Sharma R, Hanafin PO, Abboud CS, Kaye KS, et al. Evaluation Strategies for Triple-Drug Combinations against Carbapenemase-Producing Klebsiella Pneumoniae in an In Vitro Hollow-Fiber Infection Model. Clin Pharmacol Ther. 2021 Apr;109(4):1074–80.

34. Wistrand-Yuen P, Olsson A, Skarp KP, Friberg LE, Nielsen EI, Lagerbäck P, et al. Evaluation of polymyxin B in combination with 13 other antibiotics against carbapenemase-producing Klebsiella pneumoniae in time-lapse microscopy and time-kill experiments. Clin Microbiol Infect. 2020 Sep;26(9):1214–21.

35. Daikos GL, Tsaousi S, Tzouvelekis LS, Anyfantis I, Psichogiou M, Argyropoulou A, et al. Carbapenemase-Producing Klebsiella pneumoniae Bloodstream Infections: Lowering Mortality by Antibiotic Combination Schemes and the Role of Carbapenems. Antimicrob Agents Chemother. 2014 Apr;58(4):2322–8.

36. Michalopoulos A, Virtzili S, Rafailidis P, Chalevelakis G, Damala M, Falagas ME. Intravenous fosfomycin for the treatment of nosocomial infections caused by carbapenem-resistant Klebsiella pneumoniae in critically ill patients: a prospective evaluation. Clin Microbiol Infect. 2010 Feb;16(2):184–6.

37. Fosfomycin Permeation through the Outer Membrane Porin OmpF - PubMed [Internet]. [cited 2024 Nov 11]. Available from: https://pubmed.ncbi.nlm.nih.gov/30616836/

38. Fedrigo NH, Shinohara DR, Mazucheli J, Nishiyama SAB, Carrara-Marroni FE, Martins FS, et al. Pharmacodynamic evaluation of suppression of in vitro resistance in Acinetobacter baumannii strains using polymyxin B-based combination therapy. Sci Rep. 2021 May 31;11(1):11339.

39. MacNair CR, Stokes JM, Carfrae LA, Fiebig-Comyn AA, Coombes BK, Mulvey MR, et al. Overcoming mcr-1 mediated colistin resistance with colistin in combination with other antibiotics. Nat Commun. 2018 Jan 31;9(1):458.

40. Han L, Xu FM, Zhang XS, Zhang CH, Dai Y, Zhou ZY, et al. Trough polymyxin B plasma concentration is an independent risk factor for its nephrotoxicity. British Journal of Clinical Pharmacology. 2022;88(3):1202–10.

41. Imani S, Buscher H, Marriott D, Gentili S, Sandaradura I. Too much of a good thing: a retrospective study of β-lactam concentration–toxicity relationships. Journal of Antimicrobial Chemotherapy. 2017 Oct 1;72(10):2891–7.

42. Bassetti M, Peghin M, Vena A, Giacobbe DR. Treatment of Infections Due to MDR Gram-Negative Bacteria. Front Med (Lausanne). 2019;6:74.

43. Karaiskos I, Lagou S, Pontikis K, Rapti V, Poulakou G. The “Old” and the “New” Antibiotics for MDR Gram-Negative Pathogens: For Whom, When, and How. Front Public Health. 2019;7:151.

